# Two pup vocalization types are genetically and functionally separable in deer mice

**DOI:** 10.1101/2022.11.11.516230

**Authors:** N. Jourjine, M.L. Woolfolk, J.I. Sanguinetti-Scheck, J.E. Sabatini, S. McFadden, A.K. Lindholm, H.E. Hoekstra

## Abstract

Vocalization is a widespread vertebrate social behavior that is essential for fitness in the wild. While many vocal behaviors are highly conserved, heritable features of specific vocalization types can vary both within and between species, raising the questions of why and how some vocal behaviors evolve. Here, using new computational tools to automatically detect and cluster vocalizations into distinct acoustic categories, we compare pup isolation calls across neonatal development in eight taxa of deer mice (genus *Peromyscus*) and compare them to laboratory mice (C57Bl6/j strain) and free-living, wild house mice (*Mus musculus musculus*). Whereas both *Peromyscus* and *Mus* pups produce ultrasonic vocalizations (USVs), *Peromyscus* pups also produce a second call type with acoustic features, temporal rhythms, and developmental trajectories that are distinct from those of USVs. In deer mice, these tonal and low frequency “cries” are predominantly emitted in postnatal days one through nine, while USVs are primarily made after day nine. Using playback assays, we show that cries result in a more rapid approach by *Peromyscus* mothers than USVs, suggesting a role for cries in eliciting parental care early in neonatal development. Using genetic crosses between two sister species of deer mice exhibiting large, innate differences in the acoustic structure of cries and USVs, we find that variation in vocalization rate, duration, and pitch display different degrees of genetic dominance and that cry and USV features can be uncoupled in second-generation hybrids. Taken together, this work shows that vocal behavior can evolve quickly between closely related rodent species in which vocalization types, likely serving distinct functions in communication, are controlled by distinct genetic loci.

---

Vocal communication is fundamental to the social lives of vertebrates. Consistent with this critical function, vocalization is an ancient behavior, likely arising independently in multiple vertebrate lineages between 100 and 400 million years ago (Chen and Wiens, 2020; Jorgewich-Cohen et al., 2022). Since then, species have evolved differences in the acoustic features of their vocalizations and the social contexts in which those vocalizations have meaning for listeners (Jablonszky et al., 2021; Knörnschild et al., 2020; Zhao et al., 2021). In vertebrates, studies in a few exceptionally vocal groups (e.g., birds and frogs) have shed light on the ecological and social factors contributing to the evolution of this variation (Escalona Sulbarán et al., 2019; Hennelly et al., 2017; Miles et al., 2020). However, relatively little is known about the mechanisms underlying the evolution of vocal behavior, particularly among closely related species.

Recently developed computational tools have allowed for rapid, unsupervised detection of vocalizations and characterization of vocal repertoires in diverse species (e.g. Cohen et al., 2022; Fonseca et al., 2021; Odom et al., 2021; Sainburg and Gentner, 2021; Steinfath et al., 2021). These tools make it possible to measure vocal repertoires with few assumptions about what vocalizations should look like or how they should be categorized. As a result, it is now feasible to study groups of animals that, while well suited to answer questions about the evolution of vocal behavior, have received less attention from comparative studies.

One such group is the rodents. Although less well known for their vocal behaviors compared to other taxa, many rodent species are highly vocal, and they use vocalization in many of the same social contexts as other mammals (Banerjee et al., 2019; Fernández-Vargas et al., 2021; Okanoya and Screven, 2018; Pasch et al., 2013; Rieger and Marler, 2018). Studies in laboratory mice (e.g., *Mus musculus* strain C57Bl6/j) have focused on ultrasonic vocalizations (“USVs”) made by neonates (“pups”) when isolated from their parents (e.g., Zimmer et al., 2019). These pup isolation calls are thought to elicit search and retrieval behaviors from parents when pups become separated from the nest (Ehret, 2005; Ehret and Haack, 1982; Schiavo et al., 2020) and undergo a stereotyped postnatal development in their rate of production and in their acoustic features (Castellucci et al., 2018). Moreover, recent studies have begun to reveal genetic mechanisms required for pup isolation calls in *Mus* (Ashbrook et al., 2018; Barnes et al., 2016; Castellucci et al., 2016; Hernandez-Miranda et al., 2017). However, relative to laboratory strains of *Mus*, we know less about the vocal behaviors of wild *Mus* or other rodent species (but see Nicolakis et al., 2020).

Deer mice (genus *Peromyscus*) are a group of rodents that diverged from *Mus* approximately 25-40 million years ago and have since undergone a radiation across North America, resulting in several closely related, but behaviorally diverse, species (Bedford and Hoekstra, 2015). Differences have evolved between species in the vocal repertoires of both adult (Miller and Engstrom, 2012, 2007) and neonatal (Hart and King, John A., 1966) *Peromyscus* mice. And while it has been hypothesized that at least some of these differences have resulted from adaptation to specific environmental or social factors (Hart and King, John A., 1966; Kalcounis-Rueppell et al., 2018a, 2018b; Kobrina et al., 2022; Miller and Engstrom, 2012, 2007), the ultimate and proximate drivers of vocal behavior evolution in this genus remain poorly understood.

Here we focus on the postnatal development of vocal behavior in eight *Peromyscus* taxa, the C57Bl6/j strain of *Mus musculus*, and free-living, wild *Mus musculus*. Using automated detection and unsupervised clustering of vocalizations made during pup isolation assays, we find that while USVs are conserved across all these taxa, *Peromyscus* also produce low-frequency, tonal calls (“cries”). We then explore mechanisms driving variation both between call type and among species. We find that heritable vocal features have diverged quickly between *Peromyscus* species and that the distinct call types produced by *Peromyscus* pups likely serve different social functions and evolved via different genetic loci.

## Results

### *Unsupervised clustering identifies two types of pup-isolation calls in* Peromyscus

To characterize pup calls across species, we recorded isolation induced vocalizations from 577 *Peromyscus* pups belonging to four species (eight subspecies) at seven postnatal ages spanning their first two weeks of life. We compared these recordings to isolation calls from pups of the same age from laboratory *Mus musculus* (C57Bl6/j; 107 pups) and free-living, wild *Mus musculus* (109 pups) (Figure 1A; Supplemental Table 1). Using thresholding of spectrogram intensity values (Figure 1B; Supplemental Figure S1) to automatically segment these recordings into vocalizations, we first embedded spectrogram images of all vocalizations made by each taxa in two dimensions using uniform manifold approximation and projection (UMAP; Sainburg et al., 2020; Sainburg and Gentner, 2021) (Figure 1C, row 1). We found that all detected isolation calls from wild *Mus* resembled the high-frequency USVs that have been extensively characterized in C57Bl6/j, and that vocalizations from both wild and C57Bl6/j *Mus* fell into a single continuously connected cluster in UMAP space, consistent with recent descriptions of adult (Goffinet et al., 2021) and pup (Sainburg et al., 2020) C57Bl6/j vocalizations. By contrast, *Peromyscus* vocalizations separated into two distinct clusters, one of which contained short, high frequency ultrasonic frequency sweeps, while the other contained longer, tonal, low frequency vocalizations (Figure 1C, rows 2 and 3; Supplemental Figures S2 and S3).

**Figure 1.**
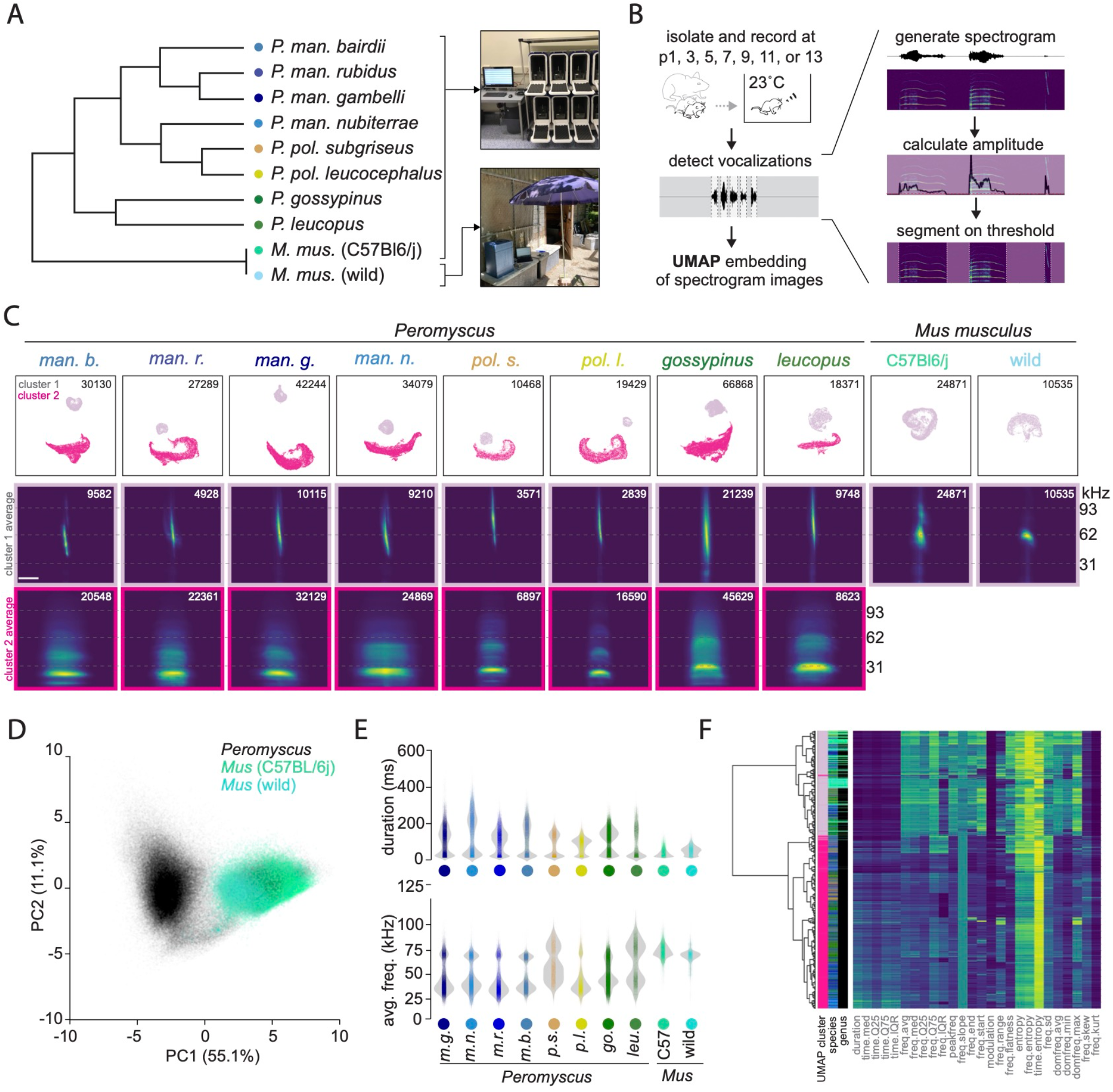
Pup isolation calls in *Peromyscus* but not *Mus* include low frequency cries. **(A)** Left: Phylogenetic relationships of taxa recorded either under laboratory conditions (top, *Peromyscus* and *Mus* C57Bl6/j) or at a field station in Zürich, Switzerland (bottom, wild *Mus musculus*). **(B)** Recording paradigm. Left: Pups were isolated and recorded at 1, 3, 5, 7, 9, 11, or 13 days after birth (day 0). Vocalizations were detected from each recording automatically using thresholding on spectrogram images. **(C)** Top: UMAP embeddings for vocalizations of each taxon, colored by HDBSCAN clustering of UMAP coordinates. Total detected vocalizations for each taxon are given (upper right-hand corner of each image). Middle and Bottom: Average spectrogram image of all vocalizations belonging to each HDBSCAN cluster for each taxon. Number of detected vocalizations in each cluster are given (upper right-hand corner). Scale bar: 100 ms. **(D)** Principal Components Analysis (PCA) on 26 acoustic features for each detected vocalization. **(E)** Top loading features from the PCA in panel D (duration and average frequency, respectively) for each taxon. **(F)** Hierarchical clustering on acoustic features of 50,000 vocalizations sampled randomly from all vocalizations. Feature names: time/freq.med = median of the energy distribution in the time/frequency domain; time/freq.Q25 = first quartile of the energy distribution in the time/frequency domain; time/freq.Q75 = third quartile of the energy distribution in the time domain; time/freq.IQR = interquartile interval of the energy distribution in the time/frequency domain.

UMAP embeddings of spectrogram images recovered meaningful variation in these vocalizations (Figure S3), but applying non-linear dimensionality reduction algorithms to spectrograms can be difficult to interpret biologically compared to more conventional bioacoustics approaches (Odom et al., 2021; Sainburg et al., 2020). We therefore calculated 26 acoustic features for each vocalization (Araya-Salas and Smith-Vidaurre, 2017) and performed principal components analysis (PCA) to measure the extent to which these features explain variation between vocalizations in the full dataset of both *Peromyscus* and *Mus*. The first two principal components (PCs) explained 55% and 10% of the variation among vocalizations, respectively, with PC1 qualitatively separating the dataset into two clusters, one of which was occupied by all taxa, while the other was occupied exclusively by vocalizations from *Peromyscus* pups (Figure 1D). The top-loading acoustic features on PC1 were duration (ms) and average frequency (kHz) of vocalizations. Plotting each of these features by taxon revealed qualitatively bi-modal distributions for all *Peromyscus*, but unimodal distributions for wild and C57Bl6/j *Mus* (Figure 1E), patterns consistent with the clustering observed in UMAP embedding of spectrogram images from these species.

We next performed hierarchical clustering of the same acoustic features calculated for each of 50,000 vocalizations sampled randomly across all recorded species, labeling the leaves of the resulting dendrogram by species and by the UMAP cluster to which each vocalizations’ spectrogram was embedded (from Figure 1B). Hierarchical clustering also split the vocalizations into two groups that corresponded to spectrogram-image based clustering: one containing short, high frequency vocalizations and one containing long, low frequency vocalizations (Figure 1F). All *Peromyscus* taxa produced vocalizations in both categories, with vocalizations from both wild and lab *Mus musculus* clustering near each other and among the short, high frequency *Peromyscus* vocalizations.

Taking these data together, automated segmentation and unsupervised clustering of 275,940 vocalizations from 10 rodent taxa suggests that isolation calls around 65 kHz are conserved between *Peromyscus* and *Mus*. We refer to this vocalization type as ultrasonic vocalizations or “USVs” as they have been referred to in previous studies of lab C57Bl6/j mice. The USVs of lab and wild house mice broadly resemble one another and cluster together in acoustic space, suggesting that laboratory domestication has not dramatically altered this behavior in *Mus*. Finally, *Peromyscus*, but not *Mus*, produce a second type of isolation call consisting of low frequency, tonal vocalizations (for examples, see Figure S2). We refer to vocalizations in this category as “cries”, as they often extend into the human hearing range and acoustically resemble the tonal cry vocalizations produced by neonates in other mammalian species.

### Interspecies variation in the low frequency cries and USVs of deer mice

We next sought to quantify interspecies variation in the acoustic features of *Peromyscus* cries and USVs. To perform analyses separately on these two vocalization types, we first annotated high quality examples of each type for each species (Supplementary Table 2). We then trained random forest classifiers to predict species identity from 14 pre-defined acoustic features for which we hypothesized interspecies variation may be relevant for pup fitness (Figure 2A). Both cry and USV classifiers predicted species above chance (12.5%), indicating that features of both vocalization types carry species-specific information. To test the robustness of cry and USV classifiers to sample size, we also trained additional random forest models on varying numbers of vocalizations, ranging from 50 to 2000. Cry and USV classifiers performed above chance with few training examples, although performance differed between species (Figure 2B, p<0.0001 for species). In addition, cry vocalizations were better predictors of species identity than USVs (Figure 2B, p<0.0001 for vocalization type, p<0.05 for (vocalization type) *(species) interaction for *P. gossypinus* and *P. maniculatus rubidus*). Together, these analyses indicate that acoustic features of both vocalization types contain information about species identity and suggest that the acoustic features of cries have diverged more among species than those of USVs.

**Figure 2.**
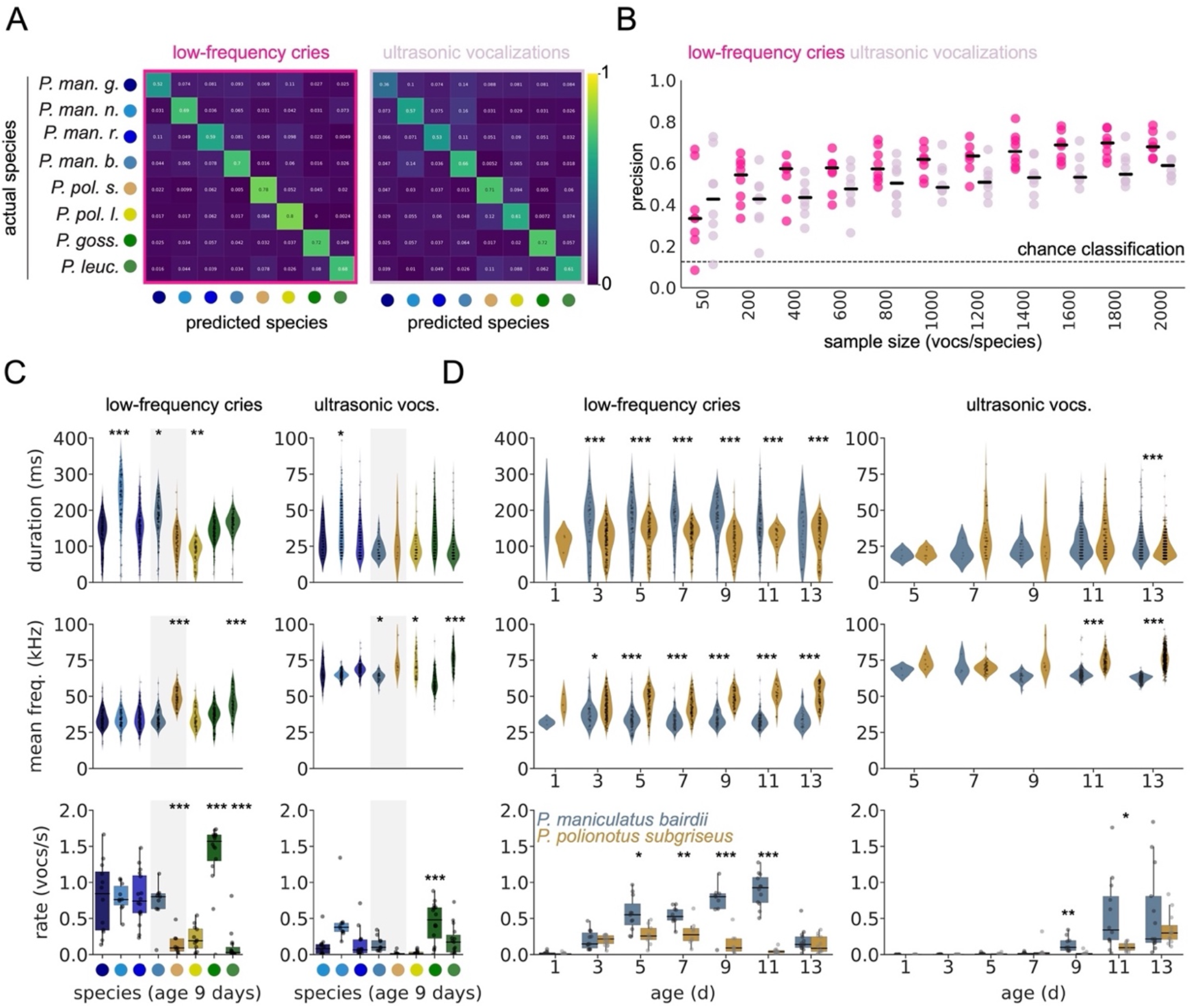
Natural variation in rate and acoustic features of *Peromyscus* pup isolation calls. **(A)** Confusion matrices showing precision of random forest classifiers (500 trees each, 80%/20% train test split) trained to predicted taxon labels from a set of hand-selected cries (left, 2000 per taxon) or USVs (right, 2000 per taxon). **(B)** Relationship between sample size and model precision of the same classifiers trained on increasing numbers of cries or USVs. Generalized linear model for statistical tests: precision ∼ (sample size)+(vocalization type)+(species)+(vocalization type)*(species). Significant effect of sample size, species, vocalization type, and species*sample size interaction (p<0.05 for interaction, p<0.0001 for others) **(C)** Duration (ms), mean frequency (kHz), and rate (vocalizations/second) of cries and USVs of p9 pups from each taxon. Gray shading highlights two focal species: *P. maniculatus bairdii* and *P. polionotus subgriseus*. Species comparison of mean frequency and duration: linear mixed effects model with species and sex as main effect and pup as random effect, with *P. maniculatus gambelli* (first column) as the reference. No effect of sex, significant effect of species (p<0.001). Species comparison of vocalization rates: ANOVA with (vocalization rate) ∼ species + sex and tukey posthoc test. No effect of sex, significant effect of species (p<0.001) **(D)** Duration (ms), frequency (kHz), and rate (vocalizations/second) of cries and USVs of *P. m. bairdii* and *P. p. subrgiseus* at all recorded ages. Species comparison of mean frequency and duration: linear mixed effects model with species, age, and species*age as main effects and pup as random effect. Species comparison of vocalization rates: ANOVA with (vocalization rate) ∼ species + age + sex + species*age and Tukey posthoc test. No effect of sex, significant effect of age, species, and species*age interaction (p<0.0001 for each). *p<0.05, **p<0.01, ***p<0.001.

To further quantify acoustic differences between cries and USVs across species, we used 3-component PCA models to reduce the dimensionality of 14 acoustic features for 500 annotated examples of each vocalization type per species (Supplementary Figure S4). Mean frequency and duration were among the top loading features of both PC1 for PC2 for both cries and USVs, suggesting they are major features that distinguish vocalizations within each of these types. Because these are also features that may be important for eliciting parental care (Lingle et al., 2012), we asked how species differed in the frequency and duration of their cries and USVs. In addition, we considered the rate and the temporal rhythms with which each species produced each vocalization type, as these have also been suggested to be functionally meaningful aspects of isolation calls (Gaub and Ehret, 2005; Lingle et al., 2012; Schiavo et al., 2020).

To make these comparisons, we first automatically labeled all detected vocalizations for each species as cry or USV using a random forest classifier (Supplementary Figure S5, out of bag score=0.996, cry F1 score=0.998, USV F1 score=0.998, non-vocal F1 score=0.985), which allowed us to analyze cries and USVs separately. We then asked how the vocalizations of *Peromyscus* species differed from one another across early postnatal development (Figure 2C, postnatal day 9 shown; Supplementary Figure S6 all ages; comparisons in 2C shown relative to the first column, *P. maniculatus gambelli*). Species differed significantly in the duration (Figure 2C top: linear mixed effects model (duration) ∼ species + sex + (1|pup), no effect of sex, p > 0.5) and mean frequency (Figure 2C middle: linear mixed effects model (mean frequency) ∼ species + sex + (1|pup), no effect of sex, p > 0.5) of their cries and USVs, as well as in the rate at which they produced each vocalization type during isolation (Figure 2C bottom: ANOVA with Tukey posthoc test, species effect p < 0.001, no effect of sex p >0.1). To compare temporal rhythms of each vocalization type, we considered the distributions of their inter-onset intervals, that is, the amount of time from the start of each vocalization to the start of the next (Burchardt and Knörnschild, 2020; Ravignani et al., 2019) (Supplementary Figure S7A). We found that the cries of all *Peromyscus* species had a similar bout-like rhythm and bi-modal distributions of inter-onset intervals (Supplementary Figure S7B, C, left), while the rhythms of USVs from all species were less clearly structured in time (Supplementary Figure S7B, C, right).

Two of the species we examined are of particular interest because they have previously been studied for their extreme differences in social system and parental care behavior: *P. maniculatus bairdii* and *P. polionotus subgriseus. P. m. bairdii* is highly promiscuous with uniparental (maternal) care of pups, while *P. p. subgriseus* is both genetically and socially monogamous, and pups receive biparental care (Bendesky et al., 2017; Birdsall and Nash, 1973; Foltz, 1981). These species differed in the duration (Figure 2D, top) and mean frequency (Figure 2D, middle) of cries and USVs as well as in the rates at which they produce them across development (Figure 2D, bottom: ANOVA with Tukey posthoc test, species effect p<0.0001, age effect p < 0.0001, species*age interactive p<0.0001, no effect of sex p>0.1). Thus, these analyses identify examples of interspecies variation in pup isolation calls between two sister species of *Peromyscus*, diverged less than ∼1 million years ago, demonstrating that these calls can evolve over short evolutionary time scales.

### Cries and USVs differ in their ability to elicit maternal approach

We find that pups from different *Peromyscus* species produce cry and USV calls at different rates across neonatal development. Cries are generally the predominant vocalization type in pups younger than nine days old, while USVs predominate in older pups (Figure 2D, bottom; Supplementary Figure S6). Since the amount of care pups require may differ between these age categories, we hypothesized that cries and USVs may signal different levels of need, and therefore would elicit responses from parents with different latencies. To test this hypothesis, we performed playback experiments in which a *P. maniculatus bairdii* mother with a nine-day old litter was presented with species-typical bouts of either cries or USVs, using recordings from nine-day old *P. m. bairdii* pups modified to be matched in amplitude and temporal rhythm (Figure 3A, B; Supplementary Figure S8). By automatically tracking the mothers’ location, we extracted spatial and velocity data and aligned these to time points when cry or USV recordings began. We found both vocalization types could cause mothers to leave their pups (Figure 3C). However, mothers were more likely to leave their pups and approach the speaker following cries (Figure 3D, E), arrived at the speaker significantly sooner after the start of vocalization playback (Figure 3F, left, paired t-test, p<0.001, mean cry=14.2s, mean USV=87.5), and reached a higher maximum velocity while moving towards the speaker (Figure 3F, right paired t-test, p<0.05, mean cry=13.5 cm/s, mean USV=6.4 cm/s). Thus, the sound of pup cries elicits more rapid behavioral responses from the mother than the sound of USVs, consistent with the hypothesis that cries are vocal signals of urgent need.

**Figure 3.**
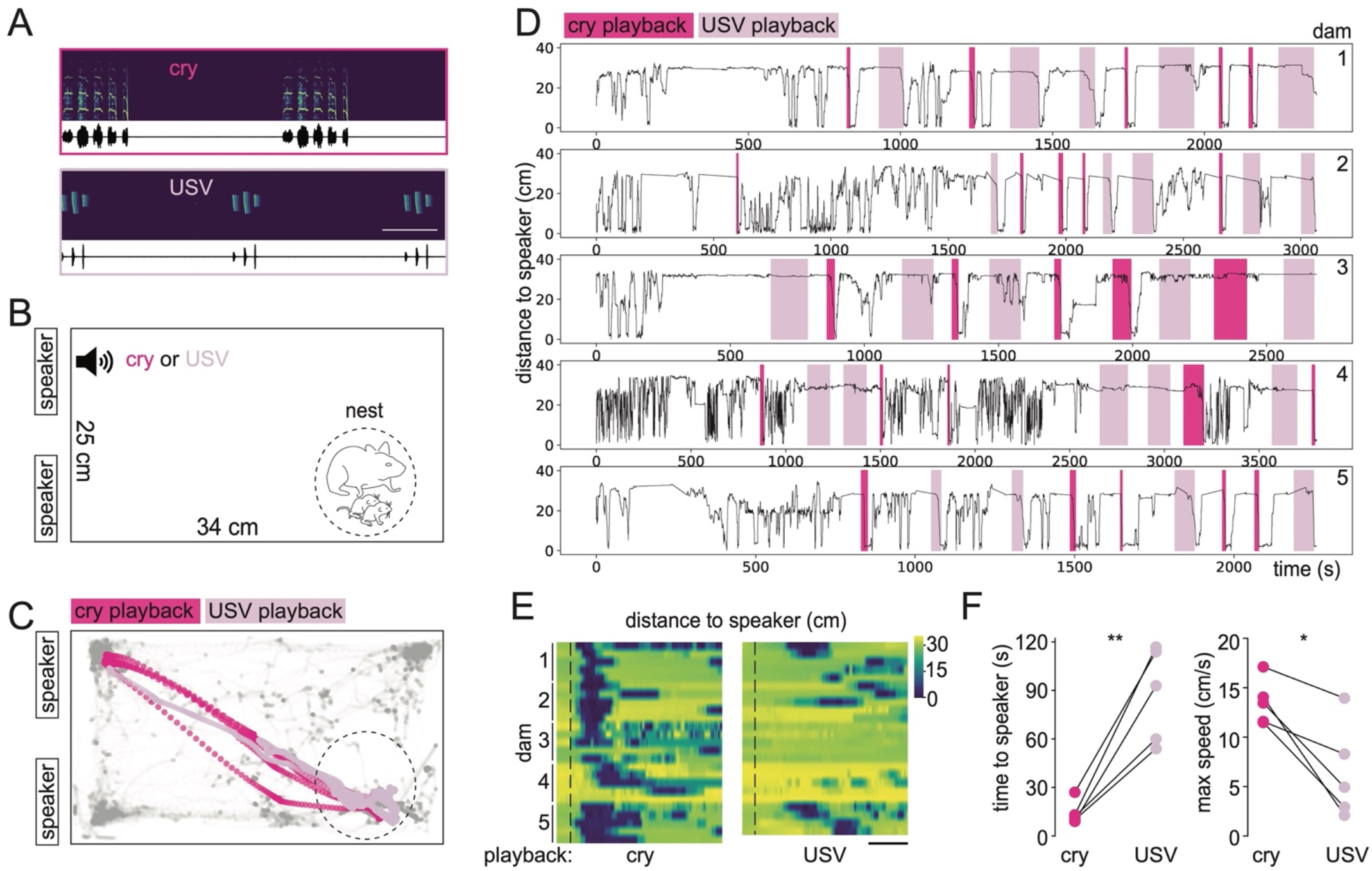
Cries elicit rapid behavior responses from *Peromyscus* mothers. **(A)** Spectrograms (top) and wave forms (bottom) of exemplar cries and USVs used during playback experiment. **(B)** Experimental setup: *P. maniculatus bairdii* mother with p9 pups was presented either the cry or USV recordings in panel A, played repeatedly in a loop from one of two speakers until either the mother reached the speaker or 2 mins elapsed, whichever came first (see Methods). Dashed circle = nest location. **(C)** Example of positional tracking (dam 5) during playback experiment. Dark pink dots = mother position in response to cries. Light pink dots = mother position in response to USVs. Grey dots = mother position with no sound. Dashed circle = nest location. **(D)** Distance of tested mothers (dam 1-5) to the active speaker. Shaded areas indicate the time periods during which cry (dark pink) or USV (light pink) stimuli were each played until the mother reached the speaker or 2 mins elapsed. **(E)** Distance to the playback speaker for each trial (row, N=5 for each mother) and mother aligned to the onset of cry (left panel) or USV (right panel). Dashed vertical line indicates start of first playback vocalization; scale bar = 30s. **(F)** Left: Median time to the speaker for each mother (N=5) following cry (dark pink) or USV (light pink) stimulus (paired t-test, **p<0.001; mean cry=14.2s, mean USV=87.5s). Right: Median of maximum speed reached for each mother following cry (dark pink) or USV (light pink) stimulus (paired t-test, *p<0.05; mean cry=13.5 cm/s, mean USV=6.4 cm/s).

### Separable genetic contributions to interspecies variation in cries and USVs

In some rodents, pup isolation calls are heritable and have been shown to respond rapidly to artificial selection (Brunelli, 2005). To identify features of deer mouse cries and USVs that have a heritable genetic contribution, we performed cross-foster experiments between the two interfertile sister species (*P. maniculatus bairdii* and *P. polionotus subgriseus*) in which we had identified differences in call rate, duration, and mean frequency (Figure 2D). When litters from both species were born on the same day, we exchanged the entire litter between parents, then recorded the pups nine days later and compared the recordings to litters from each species that were not exchanged (Figure 4A). Specifically, for each pup, we compared the median value of duration and mean frequency for all cries or USVs. We found no effect of cross fostering on the rate, mean frequency, or duration of the cries (Figure 4B, one-way ANOVA with Tukey posthoc test). We also performed PCA on 14 acoustic features calculated for the cries of each pup and found that cross-fostered pups fell into the same region of PCA space as predicted for their species, not their foster species (Figure 4C). We observed similar patterns for features of USVs with two exceptions. First, we found no significant difference in USV duration between species, and there was no effect of cross fostering on this feature. Second, the mean frequency of USVs in cross-fostered *P. p. subgriseus* were intermediate between that of non-cross-fostered pups from each species (Figure 4D, one-way ANOVA with Tukey posthoc test), suggesting that some acoustic features of USVs may be sensitive to parental environment. However, interspecific differences in USV rate were not affected by cross fostering, and PCA of USV acoustic features calculated for each pup revealed that cross-fostered pups clustered as predicted for their species and not cross-foster species, although the separation between species was smaller for USVs than for cries (Figure 4E).

**Figure 4.**
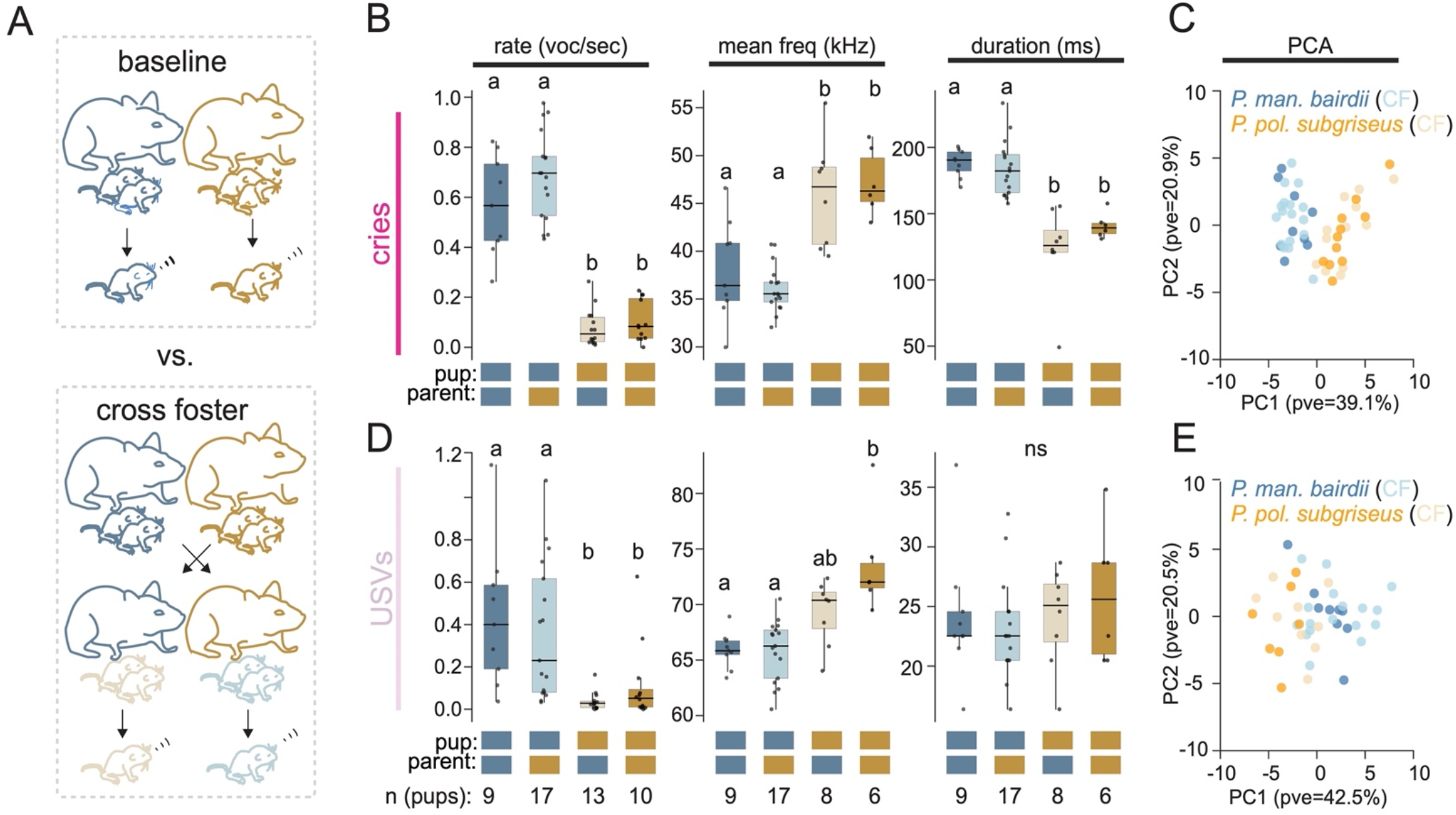
Effect of cross fostering on vocalization rate and acoustic features of cries and USVs. **(A)** Schematic of cross-fostering experimental design: pups from *P. maniculatus bairdii* (blue) and *P. polionotus subgriseus* (gold) litters born on the same day were switched within 24 h of birth and recorded in isolation at p9. **(B)** Effect of cross fostering on the rate (first column), mean frequency (second column), and duration (third column) of cries using median values or each pup. One-way ANOVA with tukey posthoc test, letters indicate significantly different groups, baseline (dark colors) and cross-fostered (light) pups. **(C)** Two-component PCA on acoustic features of cries aggregated by pup (median values for each pup). **(D)** Effect of cross fostering on the rate (first column), mean frequency (second column), and duration (third column) of USVs using median values or each pup. One-way ANOVA with tukey posthoc test, letters indicate significantly different groups, baseline (dark colors) and cross-fostered (light) pups. **(E)**Two-component PCA on acoustic features of USVs aggregated by pup (median values for each pup).

Encouraged by the large and plausibly heritable differences in vocal behavior between *P. m. bairdii* and *P. p. subgriseus*, we conducted an interspecies cross to identify genetic components that modulate vocal behavior in cries and USVs, separately. Specifically, we generated a population of first-generation (F1) hybrids and then intercrossed these F1 hybrids to generate a large population of second-generation (F2) hybrids (N= 405; Figure 5A). We found that the rate, mean frequency, and duration of low-frequency cries made by F1 hybrids all resembled those of *P. p. subgriseus* (Figure 5B, one-way ANOVA with Tukey posthoc test). PCA on all 14 acoustic features calculated for each vocalization also revealed that F1 pups occupied the same region of PCA space as *P. p. subgriseus* but not *P. m. bairdii* (Figure 5C). We observed a different pattern of inheritance for ultrasonic vocalizations. The rate of USVs made by F1 hybrids resembled that of *P. p. subgriseus*. However, the mean frequency of F1 USVs more closely resembled that of *P. m. bairdii*, and the duration of these USVs was intermediate between the two species (Figure 5D, one-way ANOVA with Tukey posthoc test). PCA revealed that F1 pups occupied a region of PCA space intermediate between *P. m. bairdii* and *P. p. subgriseus* (Figure 5E). Thus, features of cries and USVs exhibit distinct modes of inheritance in F1 hybrids, raising the possibility that cries and USVs have distinct genetic contributions.

**Figure 5.**
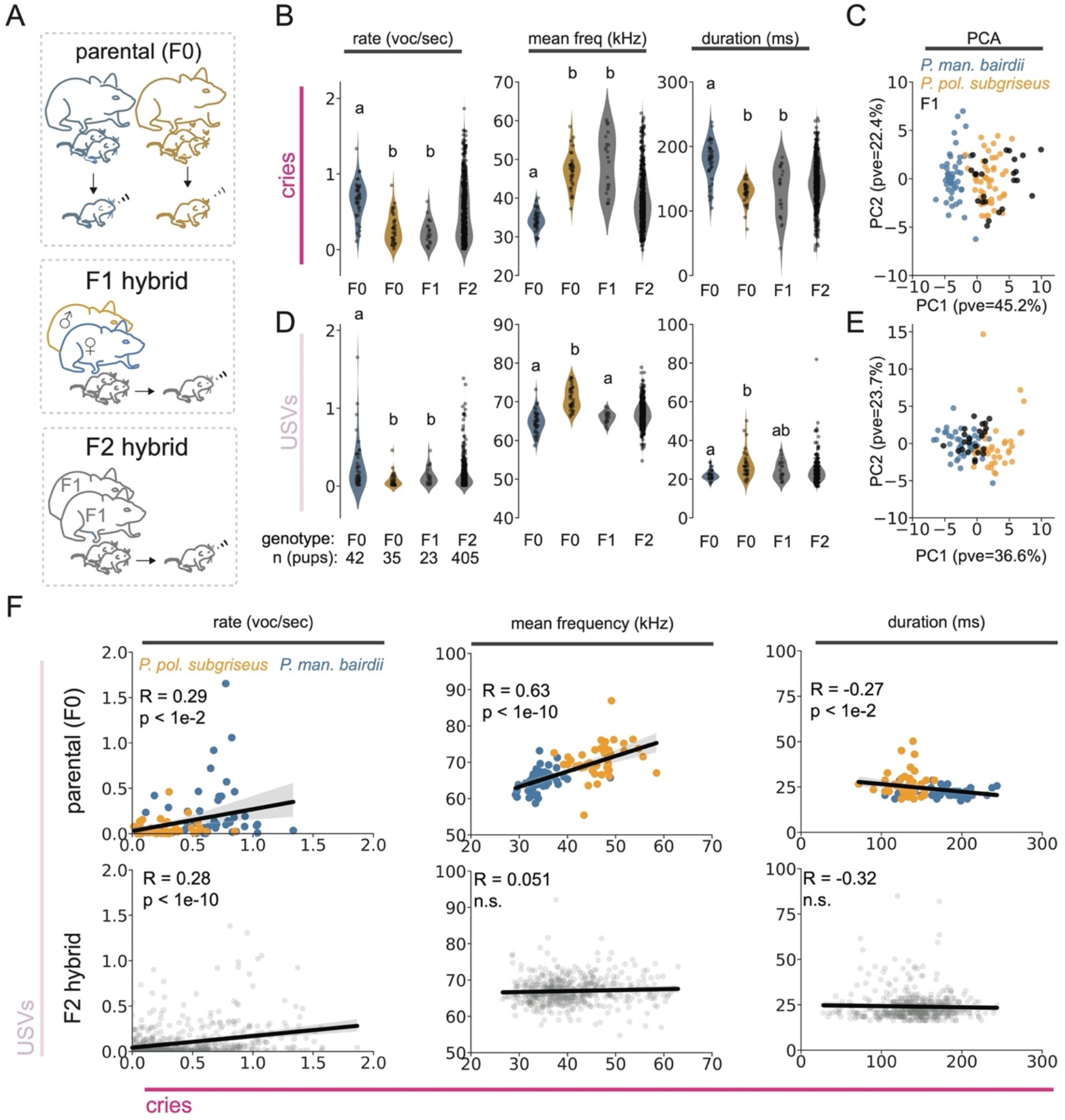
Separable genetic contributions to interspecific variation in cries and USVs. **(A)** Schematic of crossing scheme to generate first (F1) and second (F2) generation hybrids. **(B)** Comparison of rate (first column), mean frequency (second column), and duration (third column) of cries made by *P. maniculatus bairdii, P. polionotus subgriseus* and their F1 and F2 hybrids. Species and their F1 hybrids were compared by one-way ANOVA with Tukey posthoc test. Letters indicate significantly different groups (p<0.01 for all comparisons). **(C)** Two-component PCA on acoustic features of vocalizations aggregated by pup (median values for each pup). **(D)** Comparison of rate (first column), mean frequency (second column), and duration (third column) of USVs made by *P. m. bairdii, P. p. subgriseus* and their F1 and F2 hybrids. Species and their F1 hybrids were compared by one-way ANOVA with Tukey posthoc test. Letters indicate significantly different groups (p<0.01 for all comparisons). **(E)** Two-component PCA on acoustic features of USVs aggregated by pup (median values for each pup). **(F)** Correlations between cries and USVs in their rate of production (first column), mean frequency (second column), and duration (third column) in either parental species (first row) or F2 hybrids (second row). Spearman correlations were calculated for vocalization rate; Pearson correlations for mean frequency and duration.

Finally, we examined the relationship between specific features of cries and USVs in the F2 hybrids compared to correlations observed in the parental species. Vocalization rate, mean frequency, and duration were correlated in parental species (Figure 5F, top, rate: R=0.29, p < 0.01, spearman correlation, mean frequency: R=0.63, p< 0.0001, pearson correlation, duration: R = -0.27, p < 0.01, pearson correlation). To determine if variation in these vocal features of cries and USVs are genetically separable, we examined 405 F2 hybrids, each with a unique combination of alleles from the two parental species. If the same loci contribute to variation in both cries and USVs, we expect correlations between these call types to be retained in this F2 population. If different loci contribute to variation each call type, we expect the correlation between cries and USVs to be lost. We found a weak correlation in the rate at which pups produce cries and USVs in F2 hybrids (Figure 5F, bottom, R=0.28, p<0.001, spearman correlation), suggesting that the genetic loci influencing variation in the rate at which pups produce cries and USVs are partially shared. In contrast, we found no correlations between cries and USVs in their duration and mean frequency (Figure 5F, bottom, mean frequency: R = 0.051, p>0.1, pearson correlation, duration: R = -0.27, p>0.1, pearson correlation), arguing that the loci contributing to interspecies variation in these acoustic features are distinct for cries and USVs. Taken together, these data show that while interspecies variation in the rate of cries and USVs may share some genetic contribution, variation in most features, like duration and frequency, appear to be independently controlled by distinct genetic loci.

## Discussion

Using automated detection and clustering of vocal syllables, we examine the evolution of pup isolation calls in *Peromyscus*. Unlike *Mus* pups that make only USVs, we find *Peromyscus* pups produce two acoustically distinct call types: cries and USVs. These calls have distinct developmental trajectories as well as different effects on maternal behavior, with cries being produced by younger pups and triggering more rapid maternal approach behavior than USVs. By comparing pup isolation calls between two closely related species and their hybrids, we find variation in acoustic features of both cries and USVs that appear heritable, exhibit different patterns of dominance in F1 hybrids, and become largely uncoupled in F2 hybrids, suggesting that variation in vocal features can evolve rapidly via changes in distinct genetic loci.

While we identify two call types in the vocal repertoires of *Peromyscus* pups, previous studies in which vocalization types were labeled by hand have reported a larger number (e.g., “bark”, “sustained vocalization”, “simple sweep”, “complex sweep”; Kalcounis-Rueppell et al., 2018a), raising the question of how best to partition vocal repertoires into “types”. Recent unsupervised clustering of adult courtship vocalizations in C57Bl6/j *Mus* (e.g., Goffinet et al., 2021; Sainburg et al., 2020) also identified a smaller number of acoustic categories (i.e., one) than previous studies that relied on hand labeling by experts, a pattern we corroborate here for C57Bl6/j pup isolation calls. Thus, some differences between vocalizations that appear discrete to human observers are, in fact, continuous in acoustic space. In *Peromyscus*, our analyses of >250,000 isolation calls suggest that this is the case for previously described “bark” and “sustained vocalization” types, and that both fall within the “cry” category recovered by our unsupervised clustering (e.g., see Figure S2). The same is true for previously described “simple sweeps” and “complex sweeps”, which both fall within the USV type identified in this study. Importantly, the automated categorization of *Peromyscus* pup calls presented here and previously reported categorizations are each potentially informative. For example, playback experiments have shown that vocalization types with acoustic differences that are salient to humans may also be behaviorally meaningful for mice during social interactions (Pultorak et al., 2018; Rieger et al., 2021; Screven and Dent, 2019). Future work combining unsupervised clustering of vocal behaviors with playback experiments will be important for better understanding how listeners partition the acoustic space of conspecific vocalizations into meaningful categories (Sainburg and Gentner, 2021).

Unlike *Peromyscus*, isolated *Mus* pups almost exclusively vocalize in the ultrasonic range, raising the question of whether the commonly studied laboratory strains of *Mus* have a reduced vocal repertoire, perhaps due to domestication. Using a unique experimental population of wild, free-living *Mus musculus* (König et al., 2015), we find that the acoustic features of these vocalizations largely resemble those of C57Bl6/j. Like C57Bl6/j, but unlike *Peromyscus*, all wild *Mus* vocalizations are ultrasonic calls around 65 kHz, with acoustic differences between vocalizations reflecting smooth transitions (i.e., multiple small differences in acoustic features between vocalizations) rather than large discrete jumps (see Figure S2). Thus, domestication appears to have not dramatically altered the isolation calls of *Mus* pups, and the presence of cry vocalizations in *Peromyscus* but not C57Bl6/j is likely the result of evolutionary divergence in wild populations rather than an artefact of selection in the laboratory. Without a more distantly related outgroup, our study cannot definitively say which of these states (presence or absence of cries) is ancestral, although the presence of isolation cries in other rodents (e.g., lemmings, Volodin et al., 2021; fat-tailed gerbil, Zaytseva et al., 2020) and mammals more generally (Lingle et al., 2012) suggests that they were lost on the lineage leading to *Mus* rather than gained in *Peromyscus*. However, while *Mus* pups do not appear to vocalize using low-frequency sounds in pup isolation assays, pups are capable of producing audible sounds and, while less studied compared to USVs, they do so in other social contexts (e.g., “wriggling calls”; Ehret and Bernecker, 1986). Thus, the difference between *Peromyscus* and *Mus* we describe here does not reflect differences in vocal ability, but rather differences in the social contexts that elicit specific types of vocalizations.

Using playback assays, we find that cries elicit significantly faster behavioral responses from *P. maniculatus* mothers than USVs. In all the *Peromyscus* species we examined, cries are produced primarily early in postnatal development, prior to eye opening, walking, and thermoregulation, while USVs are primarily made by older pups. Thus, one hypothesis is that cries may be used when pups are most helpless and vulnerable to exposure because it garners faster attention from caregivers. This leaves open the question of the communicative function of USVs. For example, early studies in *P. maniculatus* suggested that USVs function to suppress maternal aggression (Smith, 1972, p. 197), but, to our knowledge, this hypothesis has not been tested. In *Mus*, USVs are thought to modulate maternal behavior, and the rate of USVs affects maternal responsiveness (Bowers et al., 2013; D’Amato et al., 2005; Uematsu et al., 2007).

Because USVs in *Peromyscus* do elicit maternal response (albeit more slowly than cries), one hypothesis is that, although a less salient signal, USVs may be less detectable by predators. Thus, as pups become more mature, the tradeoff between maximizing rapid parental response and minimizing detection changes. Another question is: how do species-specific differences in pup vocalizations affect maternal (or paternal) responses? In at least some mammals, mothers do not distinguish between cries of their own young and those of other species (Lingle and Riede, 2014), but as we find that acoustic features of *Peromyscus* cries have diverged between closely related and/or sympatric species, it may be important for *Peromyscus* mothers to discriminate between isolation cries of different species in the wild. Given the robust response of *Peromyscus* mothers to pup vocalizations, future playback experiments can be used to measure parental responses to interspecific variation in vocalizations as well as more fine-scale manipulation of specific acoustic features (e.g. rate, duration and frequency) of both cries and USVs.

While the function of cries versus USVs is likely distinct, the ultimate drivers of variation in these isolation calls remains unclear. One possible ecological driver is habitat. Indeed, some of the first studies of *Peromyscus* vocal behavior hypothesized that differences in vocalization rate between subspecies of *P. maniculatus* result from different selection pressures to be heard by mothers in arboreal versus terrestrial habitats (Hart and King, 1966). While some comparisons fit this hypothesis (for example, the forest dwelling species *P. maniculatus bairdii* is more vocal than its sister species, the open-field specialists *P. polionotus subgriseus and P. p. leucocephalus*), this correlation breaks down as sympatric species such as *P. maniculatus* and

*P. leucopus* vocalize at different rates. In addition, even though the four *P. maniculatus* subspecies we tested occupy different habitats, they all vocalize at similar rates. Thus, while we have not considered enough species here to perform a rigorous phylogenetic analysis, we do not observe a simple relationship between habitat use and vocalization rate.

Another possible evolutionary driver of interspecies differences is social system. In voles (genus *Microtus*), differences in pup isolation call rate have been attributed to social system complexity, with pups from monogamous, pair-bonding species vocalizing more than those from less social, promiscuous species (Blake, 2012). Four of the species we consider have well characterized differences in levels of parental care and the presence or absence of monogamy. However, in *Peromyscus*, if anything, we observe the opposite relationship between pup isolation call rate and social system to that reported in voles. For example, *P. polionotus subgriseus* is both socially and genetically monogamous, yet pups from this species are less vocal than those of the highly promiscuous *P. maniculatus bairdii*. Thus, the ultimate drivers of pup isolation call rate are likely multifaceted and may differ between species or genera in a way that belies simple correlations with habitat or social system.

We find that two pup vocalization types that are acoustically and functionally distinct have also diverged between closely related species, raising the question of how different aspects of a vocal repertoire (e.g., different call types or different acoustic features within a call type) may coevolve. For example, if different vocalization types share underlying genetic contributions, neural regulation or production mechanisms, evolutionary change in one vocalization type could result in (possibly deleterious) changes in others. On the other hand, if different vocalization types have separate underlying proximate mechanisms, changes in one call type would be less constrained, an evolutionary scenario that could facilitate functional specialization. In a comparison between two closely related species, we find that patterns of inheritance in first- and second-generation hybrids suggest that interspecies variation in vocal features of both call types are largely controlled by separate genetic loci. In comparisons of call types within species, we find that cries and USVs are produced with different temporal rhythms, suggesting at least partially distinct neural circuits pattern these behaviors (Zhang and Ghazanfar, 2020). Moreover, cries and USVs most likely arise from physically separate production mechanisms in the larynx, with the low frequency, tonal features of cries being typical of sound produced by laryngeal vocal fold vibration and high frequency sweeps of USVs likely produced by a separate mechanism (Riede et al., 2022). Taking these observations together, distinct *Peromyscus* call types appear largely unconstrained by one another, which may contribute to their functional specialization in eliciting behavioral responses from parents.

### Summary

Understanding the ultimate and proximate mechanisms driving the evolution of vocal behavior is a challenge, one that is currently being aided by rapid advances in computational tools to detect, label, and compare vocalizations across individuals and species. Using these tools in combination with playback experiments and genetic crosses, we have identified rapidly evolving features of a mammalian vocal repertoire in which interspecific variation in separable vocalization types is controlled by distinct genetic loci. Future work will identify those genetic loci and their impact on neural circuits that support social vocal communication and underlie its evolution in mammals.

## Methods

### Data Collection

#### Animal husbandry

We focused on eight *Peromyscus* taxa, representing four species (*P. maniculatus, P. polionotus, P. leucopus, P. gossypinus*). We established colonies of *P. m. bairdii, P. m. gambelli, P. p. subgriseus* and *P. leucopus* from animals originally obtained from the Peromyscus Stock Center at the University of South Carolina. We established several colonies from wild-caught animals: *P. m. nubiterrae* (Kingsley et al., 2017), *P. p. leucocephalus* (Bedford et al., 2021), *P. gossypinus* (Delaney and Hoekstra, 2018), and *P. m. rubidus* (Hager et al., 2022). We housed all animals in barrier, specific-pathogen-free conditions with 16 h light: 8 h dark at 22° C in individually ventilated cages 18.6 cm x 29.8 cm x 12.8 cm height; Allentown,

New Jersey) with quarter-inch Bed-ocob bedding (The Andersons, Maumee, Ohio). Breeding animals and their litters were fed irradiated PicoLab Mouse Diet 20 5058 (LabDiet, St. Louis, Missouri) *ad libitum* and had free access to water. We weaned animals at 23 days of age into same strain and sex cages. After weaning, we fed animals irradiated LabDiet Prolab Isopro RMH 3000 5P75 (LabDiet) *ad libitum* with free access to water and provided them with nesting material (Nestlet, Ancare, Bellmore, New York) and a polycarbonate translucent red hut. Animal experimentation protocols were approved by the Harvard University Faculty of Arts and Sciences Institutional Animal Care and Use Committee.

#### Audio recording

We recorded pups in three separate paradigms: (1) a developmental time course of *Peromyscus* and *Mus* (lab and wild) pups; (2) *P. maniculatus bairdii* and *P. polionotus subgriseus* pups that were cross-fostered; and (3) first-(F1) and second-generation (F2) hybrid pups generated from an intercross between *P. m. bairdii* and *P. p. subgriseus*. Pups were recorded in a sound-attenuating chamber (i.e., an Igloo “wheelie cool” cooler lined with acoustic foam). We collected all audio data using a multichannel recorder (Avisoft 816hb Ultrasoundgate, http://www.avisoft.com/ultrasoundgate/816h/) and Avisoft CM16/CMPA microphones (http://www.avisoft.com/ultrasound-microphones/cm16-cmpa/) at 250 kHz sampling rate and 16 bit encoding with a Windows 10 operating system.

#### (1a) *Developmental time course recordings* (*Peromyscus* and C57Bl6/J)

We recorded isolation calls from pups of eight *Peromyscus* taxa as well as an inbred strain of *Mus* (C57Bl6/j, Jackson Labs, https://www.jax.org/). We established pup age by daily nest checks, using the convention that pups are 1 day old on the day of litter discovery (day of birth is day 0). We removed breeding cages containing pups that were either 1, 3, 5, 7, 9, 11, or 13 days old from their colony room to a designated recording room and left them undisturbed for 10 mins. We then removed all pups from their nest, placed them individually into clean, empty mouse cages (18.6 cm x 29.8 cm x 12.8 cm height; Allentown, Allentown, New Jersey), and immediately recorded their temperatures using an infrared thermal camera (FLIR, model C5) directed at the back of their neck from a distance of approximately 3 inches. We then placed each pup into its own recording chamber. Then, audio recording for all pups commenced simultaneously and lasted 10 minutes, at which point we removed pups and measured their temperatures again, as described above. Pups were then weighed, sexed using ano-genital distance, and returned to the nest. Each litter/pup was recorded only once. Recordings were made in white light conditions between 5 and 2 hours prior to the transition to red light.

#### (1b) *Developmental time course recordings* (wild *Mus*)

We recorded wild *Mus musculus musculus* pups taken from a free-living population in a barn near Zürich, Switzerland that is continuously monitored as part of an ongoing, long-term study (König et al., 2015). To transfer pups, we placed pups in a clean, empty plastic container (p1 -p11) or a clean mason jar (p13, to prevent escape) and recorded each individually for 5 minutes. After audio recording, we recorded weight, sex, and age. All handling necessary for the long-term study was done following pup recording to minimize handling effects on vocal behavior. We used unique tattoo markings on pups to avoid duplicate recordings; all pups were recorded once. Recordings were made between 10am and 3pm between July 1 and 21, 2022.

#### (2) Cross-foster recordings

On days when litters were discovered (<24h old) simultaneously from *P. m. bairdii* and *P. p. subgriseus* breeding pairs, we exchanged the litters from these pairs and then returned cages to their racks until the pups were 9 days old. We found no evidence for rejection of the pups. We then removed pups from their nest into a clean Allentown cage and recorded their temperature. We recorded each pup individually, as described above, for 3 minutes. Following recording, we re-measured each pup’s temperature immediately. We then sexed, weighed, and returned each pup to their foster parents.

#### (3) F2 hybrid recordings

To generate a F2 hybrid population, we first mated two *P. m. bairdii* females to two *P. p. subgriseus* males. These founders were chosen because they had species-typical weights, heat loss, and vocalization rates when measured at p7 and p9. We then paired 54 F1 hybrids, which when paired produced F2 hybrids. All mice were paired when they were between 40 and 90 days old. We recorded from 25 F1 and 617 F2 mice, following the protocol described above. F1 and F2 hybrids were recorded twice, once at 7 days old (p7) and once at 9 days old (p9) and compared to *P. m. bairdii* and *P. p. subgriseus* pups that were also recorded twice at the same ages and under identical conditions. At p7, pups were recorded as described above for the developmental time course, with the exception that after we sexed and weighed pups, but before being returned to the nest, pups were uniquely marked on their flanks using a water-based skin marker. Pups were recorded at p9 as described above. Because we compared median values for each pup, pups that made fewer than 5 vocalizations of either type (cry or USV) were excluded from subsequent analysis.

#### Audio playback experiments

Breeding pairs of *P. m. bairdii* were checked daily for pups, which were aged as described above. When pups from a given breeding pair were 8 days old, we removed the mother, her nest, and her litter to a rat cage (25 cm x 34 cm x 19 cm height; Allentown, Allentown, New Jersey). We modified the rat cage to have two grids of holes, on either end of one wall, for audio playback. After 24 hours, we moved the cage containing the mother and her litter to a separate room and left it on a table-top for 10 minutes. Once the mother had remained in her nest for an additional 1 minute, we played pre-recorded pup vocalizations from one of the ultrasonic speakers (Avisoft, http://www.avisoft.com/playback/vifa/) until the mother touched the cage wall immediately in front of the speaker, or for 2 minutes, whichever came first. We stopped playing the audio recording until the mother returned to her nest and remained there for 1 minute, at which point we recommenced playback, using the “random” package in python to determine whether we played cries or USVs. This regime continued for 10 rounds. We used an ultrasonic microphone placed above the nest to confirm that both vocalization types were detectable at the location of the mother. The software package bonsai was used to track the mother’s position and align position measurements to the active speaker (https://open-ephys.org/bonsai). All playback experiments were performed under red light conditions. All audio was recorded using the hardware and recording specifications described above. We concatenated and modified audio clips using the free software package Audacity (https://www.audacityteam.org/). We first chose a species-typical bout of *P. m. bairdii* cries from a recording of p9 pup. We then concatenated copies this bout with alternating periods of silence. The length of this silent period was chosen to match a typical silent period between groups of cries, using data in Figure S7. A species-typical bout of USVs was chosen in the same way and concatenated with periods of silence that were matched to those of the cries. To account for the large difference in amplitude between cries and USVs and to minimize differences in background (non-vocal) silence in each recording, cry and USV recordings were matched in amplitude (USV increased to match cry) and background using the Amplify and Noise Reduction effects in Audacity.

#### Audio processing and analysis

We examined all audio during the recording using real-time spectrograms generated by Avisoft-RECORDER software.

### Segmenting the audio using amplitude thresholding

We carried out amplitude thresholding of un-preprocessed wav files in python using the get_onsets_offsets method from the python package AVA (https://github.com/pearsonlab/autoencoded-vocal-analysis). We used the following parameters for *Peromyscus* and C57Bl6/j *Mus*: Minimum frequency: 20 kHz, Maximum frequency: 125 kHz, nperseg: 1024, noverlap: 512, minimum log-spectrogram value: 0.8, maximum log-spectrogram value: 6, segmenting threshold 1: .3, segmenting threshold 2: .3, segmenting threshold 3: .35, minimum duration in seconds: 0.015, maximum duration in seconds: 1, smoothing timescale 0.00025, softmax: False. Because we recorded wild *Mus* in the field, background sound levels were higher than in laboratory recordings. We therefore segmented wild *Mus* recordings using the same parameters as above, but with the minimum log-spectrogram value of 2. For all recordings, detected vocalizations separated by less than 0.004 seconds were merged into a single vocalization.

### Acoustic feature extraction from audio segments

We calculated acoustic features from audio clips corresponding to each vocalization detected using the above parameters with the specan function of the R package warbleR (version 1.1.27, https://github.com/maRce10/warbleR; Araya-Salas and Smith-Vidaurre, 2017), with “harmonicity” set to False and “Fast” set to True.

### Unsupervised clustering of audio segments from developmental time course recordings

For each taxa recorded, we generated a spectrogram from audio clips of each detected vocalization using the following specifications: Minimum frequency: 5 kHz, Maximum Frequency 125 kHz, nperseg: 512, noverlap: 128, maximum spectrogram value: 10. To account for slight differences in background noise between recordings, we generated a background noise example for each recording that did not contain any vocalizations. For each vocalization spectrogram, we then set the minimum pixel value as 2-standard deviations above the median of the background example corresponding to the recording from which the vocalization was detected. To reduce the size of spectrogram images while preserving image features, we used the python package AVA (Goffinet et al., 2021) to pad spectrograms to length of the longest vocalization made by each species, then sampled them at 128 time points spaced evenly between the start and end of the signal (time) and 128 frequency points sampled evenly between the minimum and maximum frequency specified above (frequency), linearly interpolating between time and frequency points using the python package interp2d. This resulted in a 128×128 pixel image for each segmented vocalization. For each species, we then linearized, z-scored, and embedded these images in 2 dimensions using the default settings of the python package umap-learn (https://umap-learn.readthedocs.io/en/latest/). To label clusters of spectrogram images in these embeddings, we used the python package hdbscan (https://github.com/scikit-learn-contrib/hdbscan/blob/master/docs/index.rst) to perform unsupervised clustering of umap coordinates with the min_cluster_size set to 100, allow_single_cluster set to True, and all other parameters left as default.

### Supervised labeling of audio segments

To train random forest classifiers to label “cry” and “USV” in amplitude thresholded clips, we randomly sampled between 2000 and 6000 spectrograms from each hdbscan labeled umap cluster from each taxa using the pandas .sample method (https://pandas.pydata.org/docs/index.html). We then visually inspected each vocalization and labeled them as “cry”, “USV”, or “nonvocal sound”. We next calculated acoustic features from each of these human-verified clips as described above using the R package warbleR. The features used to described each clip were (sing warbleR naming conventions) ‘duration’, ‘time.median’, ‘time.Q25’, ‘time.Q75’, ‘time.IQR’, ‘meanfreq’, ‘freq.median’, ‘freq.Q25’, ‘freq.Q75’, ‘freq.IQR’, ‘meanpeakf’, ‘dfslope’, ‘enddom’, ‘startdom’, ‘modindx’, ‘dfrange’, ‘sfm’, ‘entropy’, ‘sp.ent’, ‘time.ent’, ‘sd’, ‘meandom’, ‘mindom’, ‘maxdom’, ‘skew’, and ‘kurt’. We then used vestors of these features corresponding labels for each sampled vocalization to train a 10000 tree Random Forest Classifier using the RandomForestClassifier class from the python package sklearn (https://scikit-learn.org/stable/) and an 80%/20% train/test split. Information gain was used as the optimization criterion (the ‘criterion’ parameter was set as ‘entropy’).

### Predicting species from acoustic features of cry and ultrasonic vocalizations

To assess the species-specificity of cry and USV acoustic features, we trained 500 tree random forest classifiers on of human-validated cries or USVs consisting of either 50, 200, 400, 600, 800, 1000, 1200, 1400, 1600, 1800, or 2000 vocalizations per call type, sampled using the pandas .sample method, as above. For each sample size and vocalization type (cry or USV), we then used the RandomForestClassifier class from the python package sklearn and an 80%/20% train/test split, as above. Information gain was used as the optimization criterion (the ‘criterion’ parameter was set as ‘entropy’).

## Respective contributions

NJ and HEH conceived and planned the project. NJ, MLW, JIS, JES, and SM collected the data. NJ, MLW, JIS, and JES analyzed the data. AKL provided access to wild *Mus musculus* and input on interpreting recordings of these mice. NJ and HEH wrote the manuscript with feedback from all co-authors.

## Animal Experimentation

All experiments on *Peromyscus* and *Mus* (C57Bl6/j) were approved by the Harvard University IACUC. All recordings of wild *Mus musculus* were approved under permit ZH076/2022 granted by the Veterinary Office of Canton Zürich, Switzerland.

## Data Availability

All code used for data analysis will be made publicly available on GitHub upon publication. All data will be made available in a public repository upon publication.

## Acknowledgements

We would like to thank Barbara König and the members of the Hoekstra and Lindholm labs for their feedback and advice throughout the course of this research, Andri Manser for advice on statistical analyses, and the staff of the Harvard University Animal Facility for care of *Peromyscus* and C57Bl6/j mouse colonies. We also thank Tim Sainburg, Gregg Castellucci, and Kelsey Tyssowski for their thoughtful comments on the manuscript.

## Supplemental Tables and Figures

**Table S1 (related to Figure 1).** Pup, litter, and vocalization counts by species.

| species | # pups recorded (all ages) | # detected vocalizations | # litters recorded per age (age with each number of litters) |
| --- | --- | --- | --- |
| *P. maniculatus bairdii* | 76 | 30,130 | 4 (p3); 3 (all other ages) |
| *P. maniculatus rubidus* | 73 | 27,289 | 2 (p1,p5,p7); 3 (all other ages) |
| *P. maniculatus gambelli* | 97 | 42,244 | 3 (all ages) |
| *P. maniculatus nubiterrae* | 70 | 34,079 | 3 (all ages) |
| *P. polionotus subgriseus* | 71 | 104,68 | 3 (all ages) |
| *P. polionotus leucocephalus* | 66 | 18,371 | 3 (all ages) |
| *P. gossypinus* | 66 | 66,868 | 5 (p9); 3 (all other ages) |
| *P. leucopus* | 58 | 18,371 | 2 (p1, p13); 6 (p9); 3 (all other ages) |
| *Mus musculus* (C57Bl6/j) | 107 | 26,543 | 2 (p1, p5, p9, p13); 3 (all other ages) |
| *Mus musculus* (Wild) | 109 | 12,045 | 2 (p11); 4 (p3, p7, p13); 5 (all other ages) |
| **Total** | 793 | 275,940 | 213 |

**Table S2 (related to Figure 2).** Number of human annotated vocalizations by vocalization type and species.

| species | # cries annotated | # USVs annotated | **Total** |
| --- | --- | --- | --- |
| *P. maniculatus bairdii* | 4,734 | 13,730 | 18,464 |
| *P. maniculatus rubidus* | 4,824 | 8,608 | 13,432 |
| *P. maniculatus gambelli* | 5,194 | 10,574 | 15,768 |
| *P. maniculatus nubiterrae* | 4,542 | 9,832 | 14,374 |
| *P. polionotus subgriseus* | 5,198 | 6,130 | 11,328 |
| *P. polionotus leucocephalus* | 4,076 | 4,678 | 8,754 |
| *P. gossypinus* | 4,678 | 13,320 | 17,998 |
| *P. leucopus* | 4,184 | 12,048 | 16,232 |
| **Total** | 37,430 | 78,920 | 116,350 |

**Figure S1 (related to Figure 1).**
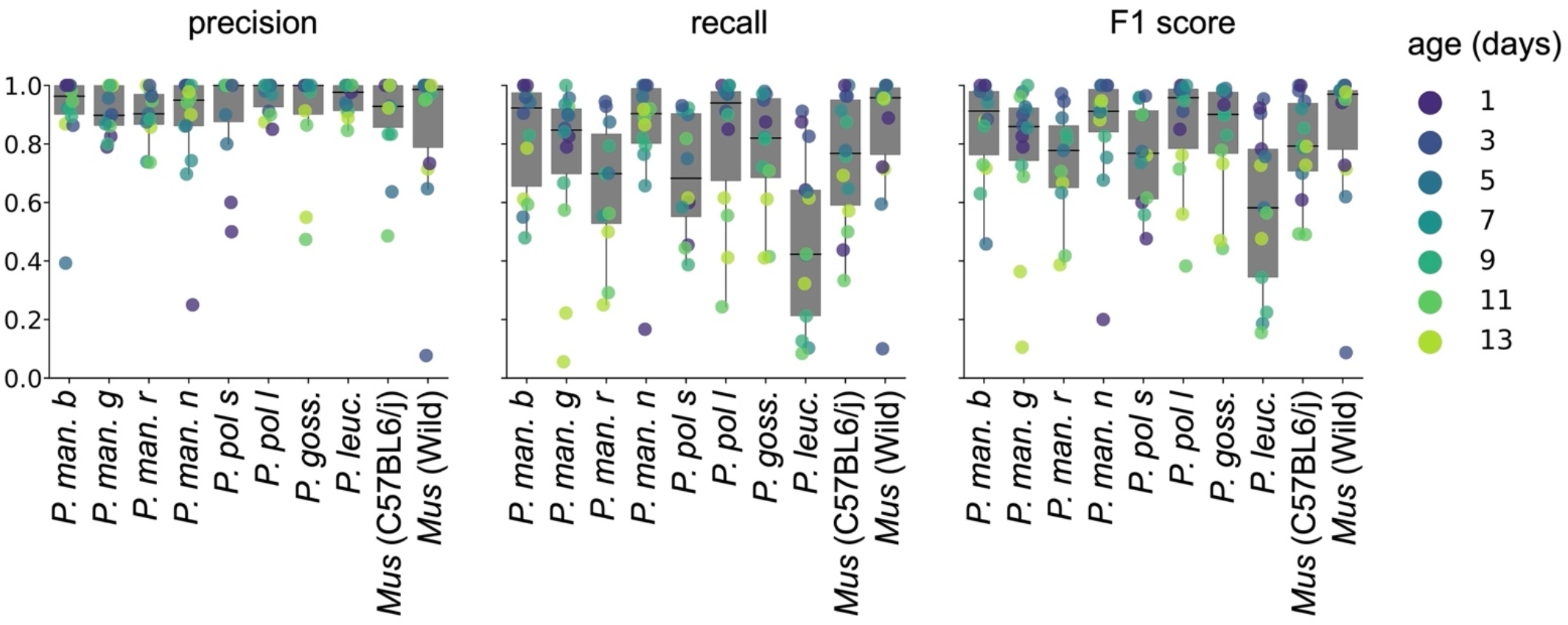
Evaluation of segmenting algorithm. Segmenting parameters were evaluated on 30 sec recordings from 2 pups from each age and each taxon (112 total recordings with varying numbers of vocalizations). For each vocalization, if the predicted start and stop were each within a 20 ms window of the true start and stop, the prediction was considered a true positive. Left: precision (i.e., percent of predicted vocalizations that were true vocalizations for each recording). Middle: recall (i.e., percent of true vocalizations that were detected for each recording). Right: F1 score, calculated as 2 * (precision * recall) / (precision + recall).

**Figure S2 (related to Figure 1).**
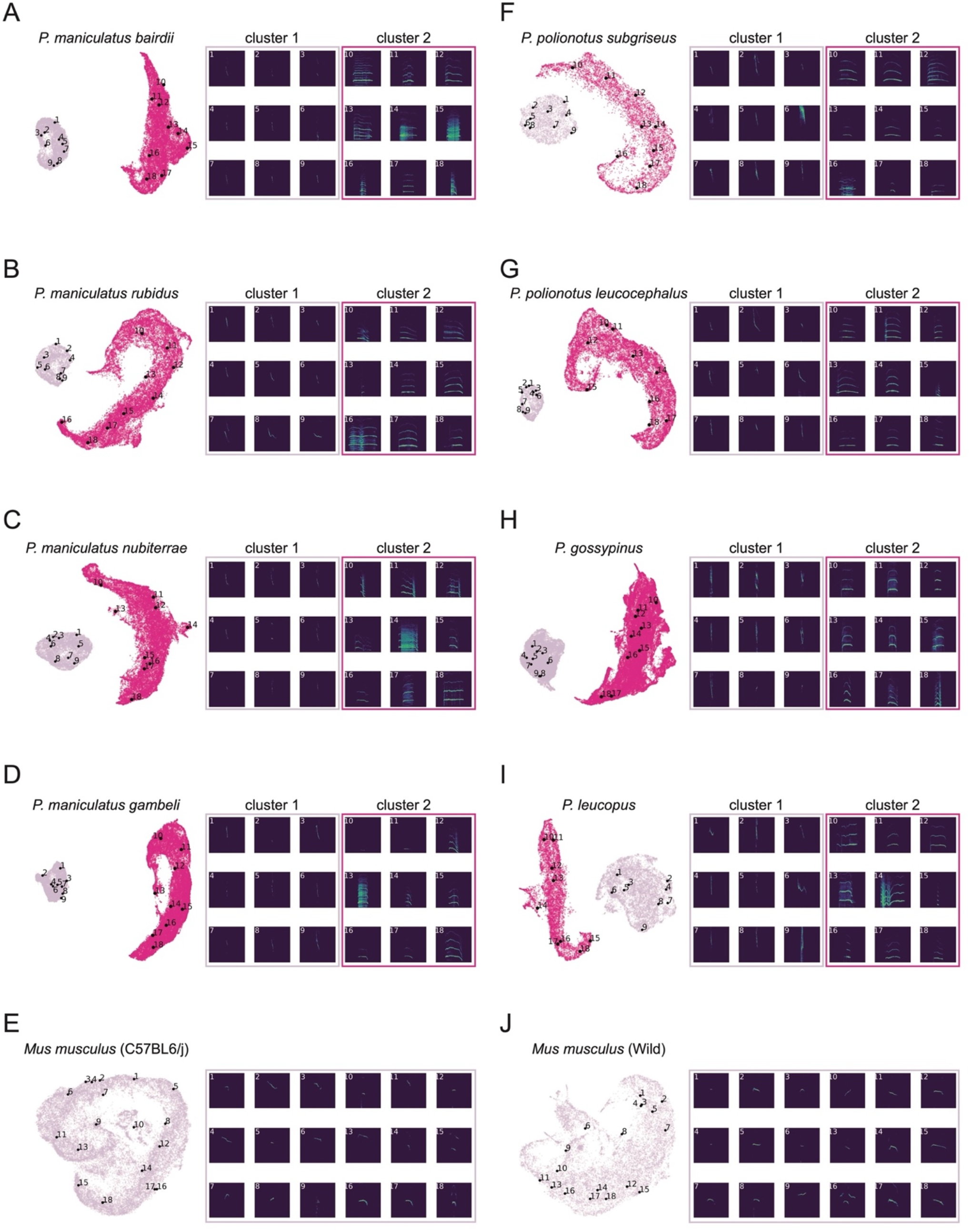
Example spectrogram images from UMAP embeddings. For each taxon (A-J), nine vocalizations were randomly sampled from within each unsupervised clustering label (cluster 1 or cluster 2 for *Peromyscus*; no clusters for *Mus*). For each taxon, vocalization location in UMAP is space shown and labeled with a number (1-18) on the left. Corresponding labeled spectrogram image are shown on the right.

**Figure S3 (related to Figure 1).**
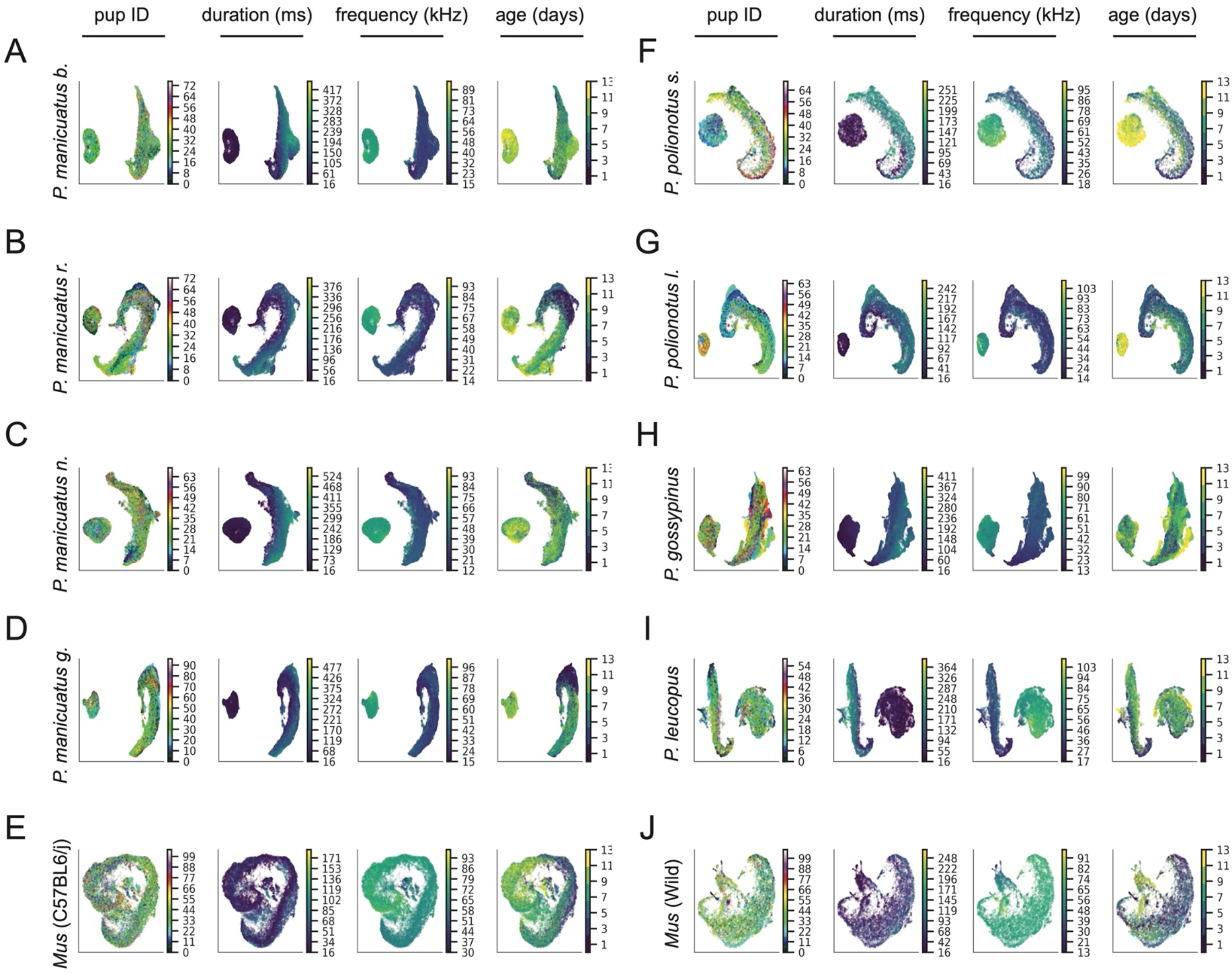
Correspondence between UMAP embeddings and acoustic features. UMAP embeddings of all detected vocalizations colored by pup ID, duration (ms), mean frequency (kHz), and age (in days, day of birth is day 0) for each taxon (A-J).

**Figure S4 (related to Figure 2).**
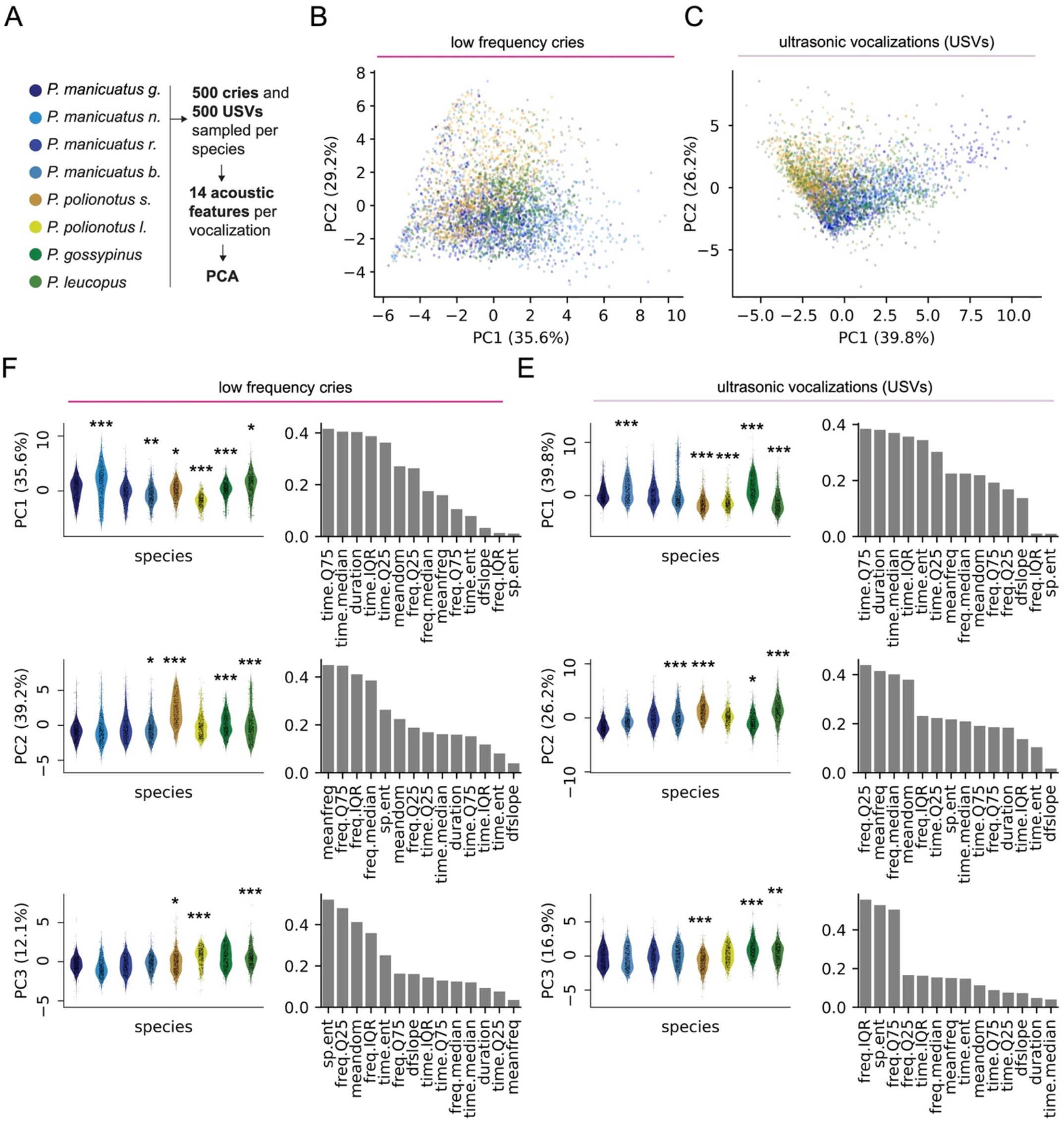
Principal components analysis on acoustic features of cries and USVs. **(A)** Schematic of sampling scheme: 500 cries and 500 USVs were sampled per *Peromyscus* taxon. 14 acoustic features were calculated for each vocalization, and PCA was then run on these features either for all cries or all USVs. **(B)** Scatter plot of the first two PCs from a three-component model for cries (500 per species sampled across all ages). Dot color corresponds to taxon. **(C)** Scatter plot of the first two PCs from a three-component model for USVs (500 per species sampled across all ages). Dot color corresponds to taxon. **(D)** Plot of PCs by taxon (left) and feature loadings on each PC axis (right) for cries. Plot of PCs by taxon (left) and feature loadings on each PC axis (right) for USVs. *p<0.05, **p<0.01, ***p<0.0001. PC values were compared across taxa with linear mixed effects models with species and sex as main effects and pup as a random effect. Significant effect of species (p < 0.01 for all comparisons), no effect of sex. Reference for comparisons is *P. maniculatus gambelli* (first species on the left in all plots).

**Figure S5 (related to Figure 2).**
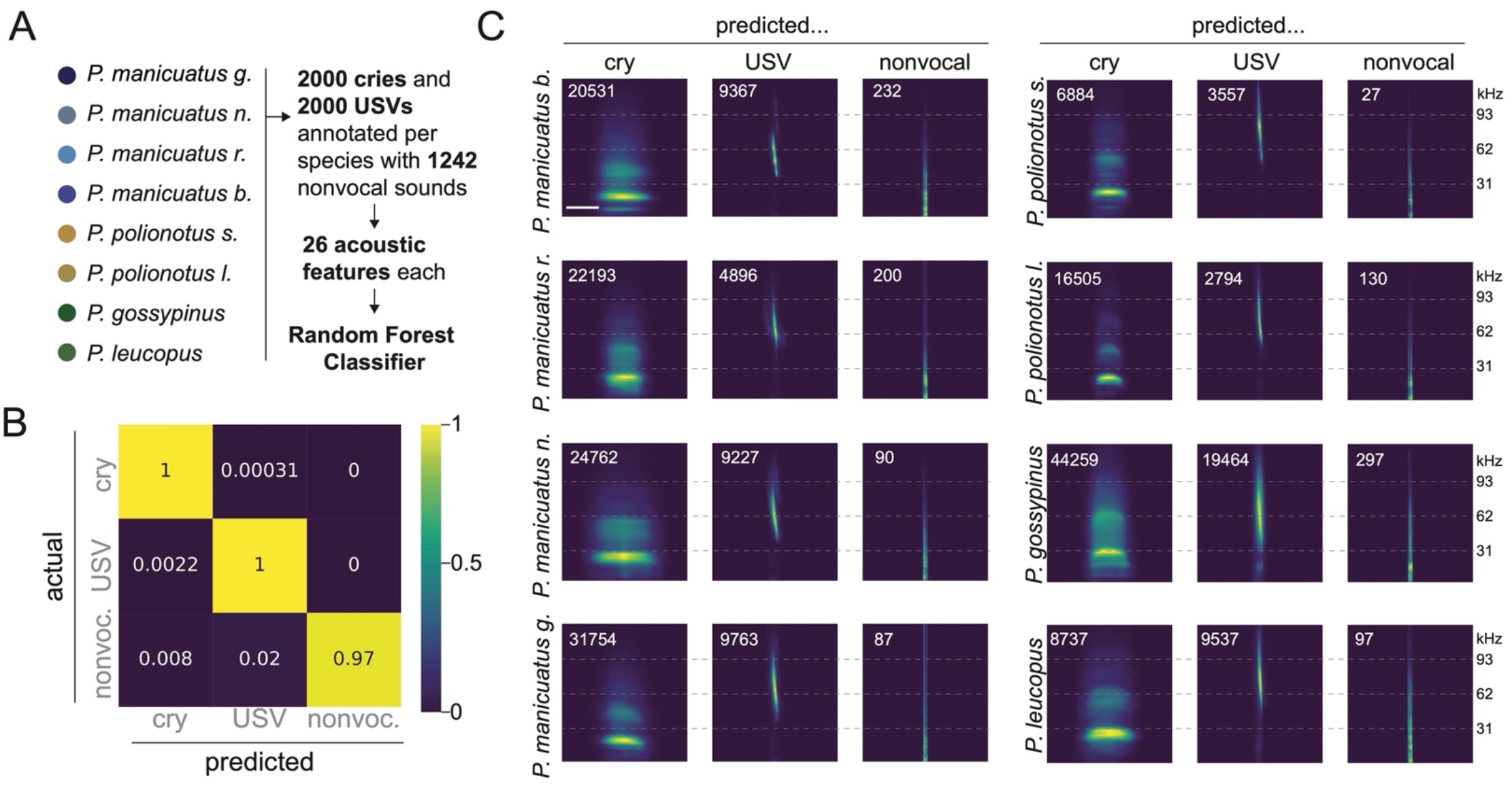
Evaluation of machine learning classifier to label amplitude segmented vocalization as “cry” or “USV”. **(A)** Schematic of the training data and model. 2000 cries and 2000 USVs were annotated by hand for each taxon. A total of 1,242 nonvocal sounds were also annotated across all taxa. 26 acoustic features were calculated for each sound, and these features were used to train a random forest classifier on the vocalization type labels “cry”, “USV”, and “nonvocal” (“nonvoc.”) (10,000 trees, entropy as the optimization metric, 80%/20% train/test split). **(B)** Confusion matrix showing performance of the classifier on test data. **(C)** Average spectrograms of all predicted cries, USVs and nonvocal sounds, for all detected sounds from all taxa. Number of vocalizations in the average in the upper right corner of each image. Scale bar: 100 ms.

**Figure S6 (related to Figure 2).**
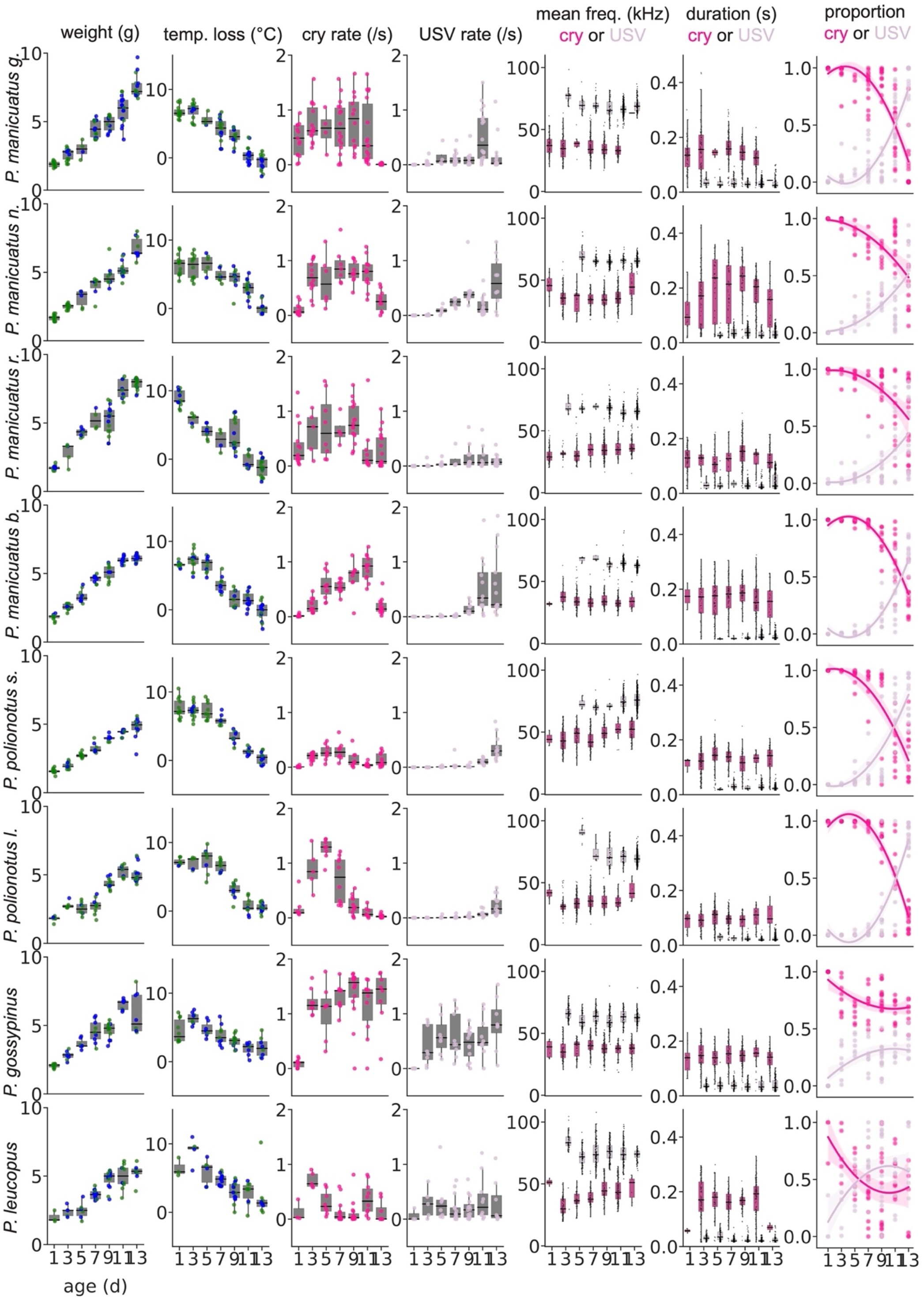
Vocal and non-vocal development of *Peromyscus* pups between ages p1 and p13. From left to right: pup weight at each age recorded for each taxon (male: green, female: blue); amount of heat lost by pup during recording (°C, calculated ass (−1)*((temperature post recording) – (temperature pre recording)); cry vocalization rate, USV vocalization rate, mean frequency (Hz), duration (s), the proportion of all vocalizations that were either cry or USV was calculated. Proportions for each vocalization type were fit with a second order polynomial (dark line). Shading above and below lines are 95% confidence intervals.

**Figure S7 (related to Figure 2).**
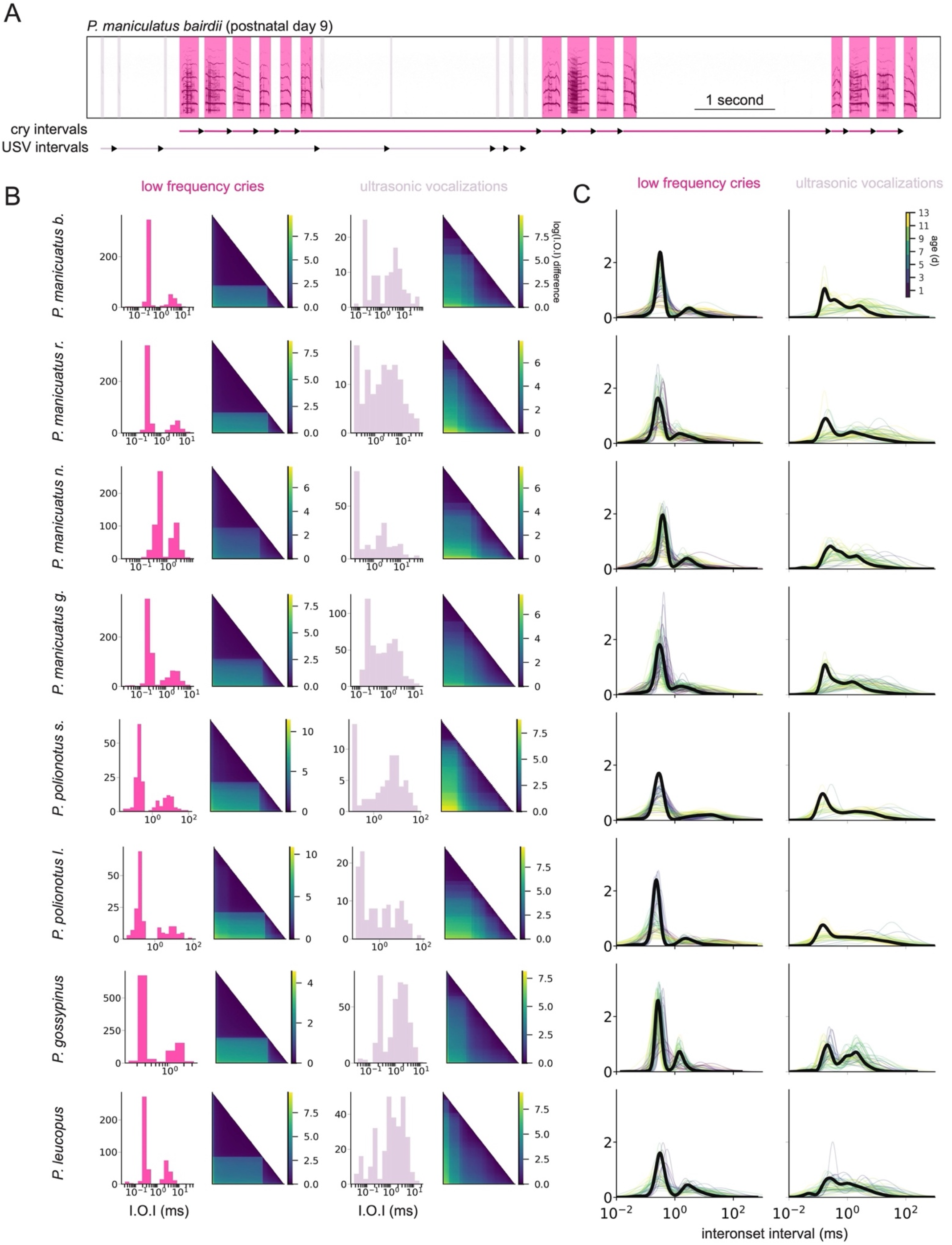
Different temporal rhythms between cries and USVs within taxa. **(A)** Example spectrogram from a p9 *P. maniculatus bairdii* pup with inter-onset intervals of cries (“cry intervals”) or USVs (“USV intervals”) labeled on the time axis. Black arrow tips indicate the end of each inter-onset interval. **(B)** Histograms (left) and recurrence plots (right) of all inter-onset intervals (I.O.I) for cries and USVs made by one example pup per taxon. Recurrence plots were generated by taking the logarithm of all inter-onset intervals of a given vocalization type, the difference of every log(interonset interval) with every other log(interonset interval), then sorting the differences from smallest to largest and plotting as a heatmap. **(C)** Smoothed histograms of all inter-onset intervals of cries (left) and USVs (right). Thin lines: individual pups colored by their age. Dark black line: all pups pooled.

**Figure S8 (related to Figure 3).**
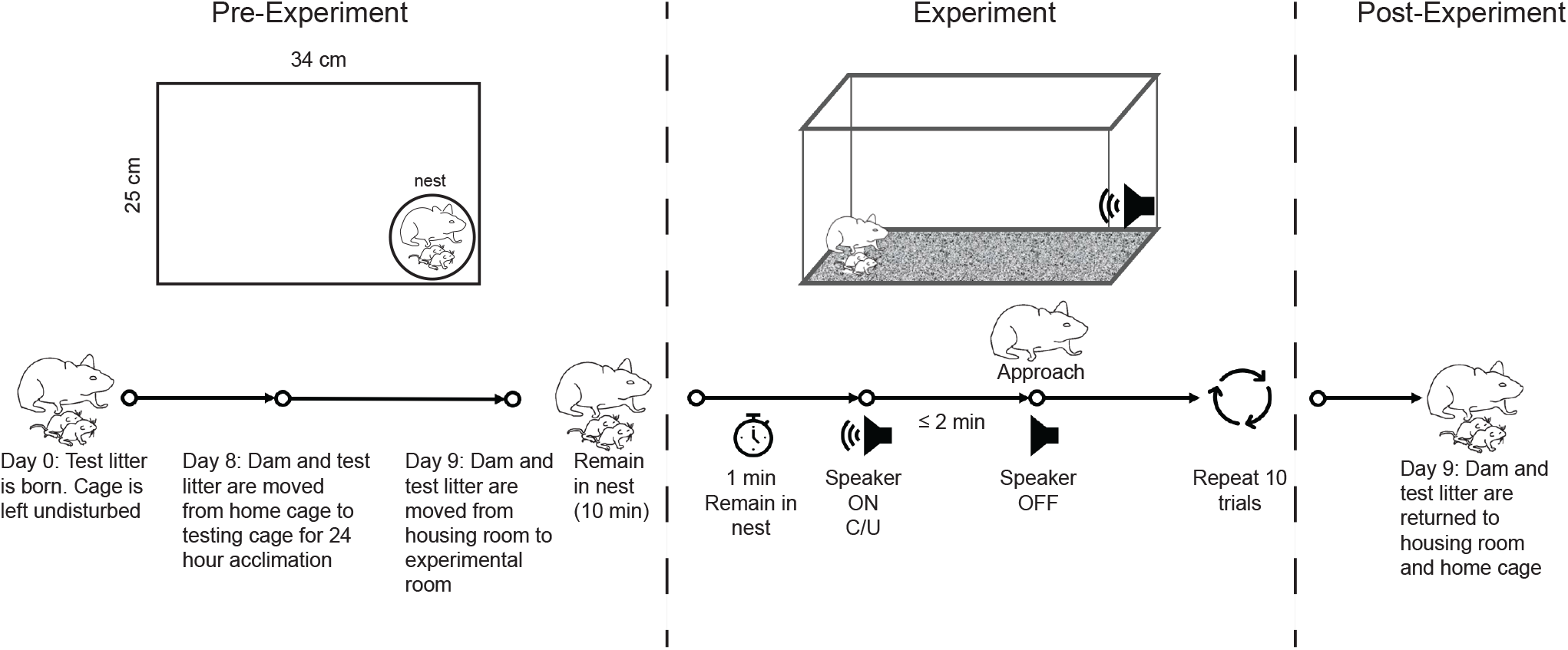
Experimental protocol for playback experiments. Pre-experiment: Mothers and their pups were acclimated for 24 h in a large cage outfitted with 2 speakers. Experiment: We played recordings of cries or USVs in a randomized order from a single speaker in the opposite corner from the nest. Prior to each trial, mothers were required to remain in the nest continuously for 1 min. The audio recordings were played until the mother reached the speaker or 2 mins elapsed. Post-experiment: All animals were returned to their home cage. C: cry; U: USV.

